# Loss-of-function genomic variants with impact on liver-related blood traits highlight potential therapeutic targets for cardiovascular disease

**DOI:** 10.1101/597377

**Authors:** Jonas B. Nielsen, Oren Rom, Ida Surakka, Sarah E. Graham, Wei Zhou, Lars G. Fritsche, Sarah A. Gagliano Taliun, Carlo Sidore, Yuhao Liu, Maiken E. Gabrielsen, Anne Heidi Skogholt, Brooke Wolford, William Overton, Whitney E. Hornsby, Akua Acheampong, Austen Grooms, Tanmoy Roychowdhury, Amanda Schaefer, Gregory JM Zajac, Luis Villacorta, Jifeng Zhang, Ben Brumpton, Mari Løset, Vivek Rai, Kent D. Taylor, Nicholette D. Palmer, Yii-Der Chen, Seung Hoan Choi, Steven A. Lubitz, Patrick T. Ellinor, Kathleen C. Barnes, Michelle Daya, Nicholas Rafaels, Scott T. Weiss, Jessica Lasky-Su, Russell P. Tracy, Ramachandran S. Vasan, L. Adrienne Cupples, Rasika A. Mathias, Lisa R. Yanek, Lewis C. Becker, Patricia A. Peyser, Lawrence F. Bielak, Jennifer A. Smith, Stella Aslibekyan, Bertha A. Hildalgo, Donna K. Arnett, Marguerite R. Irvin, James G. Wilson, Solomon K. Musani, Adolfo Correa, Stephen S. Rich, Xiuqing Guo, Jerome I. Rotter, Barbara A. Konkle, Jill M. Johnsen, Allison E. Ashley-Koch, Marilyn J. Telen, Vivien A. Sheehan, John Blangero, Joanne E. Curran, Juan M. Peralta, Courtney Montgomery, Wayne H-H Sheu, Ren-Hua Chung, Karen Schwander, Seyed M. Nouraie, Victor R. Gordeuk, Yingze Zhang, Charles Kooperberg, Alexander P. Reiner, Rebecca D. Jackson, Eugene R. Bleecker, Deborah A. Meyers, Xingnan Li, Sayantan Das, Ketian Yu, Jonathon LeFaive, Albert Smith, Tom Blackwell, Daniel Taliun, Sebastian Zollner, Lukas Forer, Sebastian Schoenherr, Christian Fuchsberger, Anita Pandit, Matthew Zawistowski, Sachin Kheterpal, Chad M. Brummett, Pradeep Natarajan, David Schlessinger, Seunggeun Lee, Hyun Min Kang, Francesco Cucca, Oddgeir L. Holmen, Bjørn O. Åsvold, Michael Boehnke, Sekar Kathiresan, Goncalo Abecasis, Y. Eugene Chen, Cristen J. Willer, Kristian Hveem

**Affiliations:** Department of Internal Medicine: Cardiology, University of Michigan, Ann Arbor, Michigan, USA; Program in Medical and Population Genetics, Broad Institute of Harvard and MIT, Cambridge, Massachusetts, USA; Analytic and Translational Genetics Unit, Massachusetts General Hospital, Boston, Massachusetts, USA; Stanley Center for Psychiatric Research, Broad Institute of Harvard and MIT, Cambridge, Massachusetts, USA; Department of Computational Medicine and Bioinformatics, University of Michigan, Ann Arbor, MI, USA; Department of Biostatistics, University of Michigan School of Public Health, Ann Arbor, Michigan, USA; Center for Statistical Genetics, University of Michigan School of Public Health, Ann Arbor, Michigan, USA; Istituto di Ricerca Genetica e Biomedica, Consiglio Nazionale delle Ricerche (CNR), Monserrato, Cagliari, Italy; K.G. Jebsen Center for Genetic Epidemiology, Department of Public Health and Nursing, Faculty of Medicine and Health Sciences, Norwegian University of Science and Technology, NTNU, Norway; Department of Dermatology, St. Olav’s Hospital, Trondheim University Hospital, Trondheim, Norway; Department of Human Genetics, University of Michigan, Ann Arbor, Michigan, USA; The Institute for Translational Genomics and Population Sciences, Department of Pediatrics and Los Angeles Biomedical Research Institute, Harbor-UCLA, Torrance, California, USA; Department of Biochemistry, Wake Forest School of Medicine, Winston-Salem, North Carolina, USA; Cardiovascular Research Center, Massachusetts General Hospital, Boston, Massachusetts, USA; Colorado Center for Personalized Medicine, School of Medicine, University of Colorado, Aurora, Colorado, USA; Channing Division of Network Medicine, Department of Medicine Brigham and Women’s Hospital, Boston, Massachusetts, USA; Harvard Medical School, Boston, Massachusetts, USA; Department of Pathology and Laboratory Medicine, Larner College of Medicine, University of Vermont, Burlington, Vermont, USA; Department of Biochemistry, Larner College of Medicine, University of Vermont, Burlington, Vermont, USA; Department of Medicine, Boston University School of Medicine, Boston, MA 02118, USA; Framingham Heart Study, Framingham, Massachusetts, USA; Department of Biostatistics, Boston University School of Public Health, Boston, Massachusetts, USA; GeneSTAR Research Program, Department of Medicine, Johns Hopkins School of Medicine, Baltimore, Maryland, USA; Department of Epidemiology, School of Public Health, University of Michigan, Ann Arbor, Michigan, USA; Survey Research Center, Institute for Social Research, University of Michigan, Ann Arbor, Michigan, USA; The University of Alabama at Birmingham, Birmingham, Alabama, USA; 23andMe, Inc; Deans Office, College of Public Health, University of Kentucky, Lexington, Kentucky, USA; Department of Physiology and Biophysics, University of Mississippi Medical Center, Jackson, Mississippi, USA; Jackson Heart Study, Jackson, Mississippi, USA; Department of Medicine, University of Mississippi Medical Center, Jackson, Mississippi, USA; Center for Public Health Genomics, University of Virginia, Charlottesville, Virginia, USA; BloodWorks Northwest, University of Washington, Seattle, Washington, USA; Duke Molecular Physiology Institute, Duke University Medical Center, Durham, North Carolina, USA; Department of Medicine, Duke University Medical Center, Durham, North Carolina, USA; Department of Pediatrics, Division of Hematology/Oncology, Baylor College of Medicine, Houston, Texas, USA; Department of Human Genetics and South Texas Diabetes and Obesity Institute, University of Texas Rio Grande Valley School of Medicine, Brownsville Texas, USA; Department of Genes and Human Disease, Oklahoma Medical Research Foundation, Oklahoma, USA; Division of Endocrinology and Metabolism, Department of Internal Medicine, Taichung Veterans General Hospital, Taichung, Taiwan; Institute of Population Health Sciences, National Health Research Institutes, Miaoli, Taiwan; Division of Biostatistics, Washington University School of Medicine, St. Louis, Missouri, USA; University of Pittsburgh School of Medicine, Pittsburgh, Pennsylvania, USA; University of Illinois at Chicago, Chicago, Illinois, USA; Division of Public Health Sciences, Fred Hutchinson Cancer Research Center, Seattle, Washington, USA; Department of Epidemiology, University of Washington, Seattle, Washington, USA; Division of Endocrinology, Diabetes and Metabolism, Ohio State University, Columbus, Ohio, USA; Division of Pharmacogenomics University of Arizona, Tucson, Arizona, USA; Division of Genetics, Genomics and Precision Medicine, Department of Medicine, University of Arizona, Tucson, Arizona, USA; Division of Genetic Epidemiology, Department of Medical Genetics, Molecular and Clinical Pharmacology, Medical University of Innsbruck, Innsbruck, Austria; Institute for Biomedicine, Eurac Research, Bolzano, Italy; Department of Anesthesiology, University of Michigan, Ann Arbor, Michigan, USA; Center for Genomic Medicine and Cardiovascular Research Center, Massachusetts General Hospital, Boston, Massachusetts, USA; Laboratory of Genetics, National Institute on Aging, US National Institutes of Health, Baltimore, Maryland, USA; Dipartimento di Scienze Biomediche, Università degli Studi di Sassari, Sassari, Italy; HUNT Research Centre, Department of Public Health and Nursing, Norwegian University of Science and Technology, Levanger, Norway; Department of Endocrinology, St. Olavs Hospital, Trondheim University Hospital, Trondheim, Norway; Broad Institute, Cambridge, Maryland, USA; Regeneron Pharmaceuticals, Tarrytown, New York, USA.USA

**Keywords:** genome-wide, phenome-wide, liver, lipids, iron, coding.

## Abstract

Cardiovascular diseases (CVD), and in particular cerebrovascular and ischemic heart diseases, are leading causes of death globally.^1^ Lowering circulating lipids is an important treatment strategy to reduce risk.^2,3^ However, some pharmaceutical mechanisms of reducing CVD may increase risk of fatty liver disease or other metabolic disorders.^4,5,6^ To identify potential novel therapeutic targets, which may reduce risk of CVD without increasing risk of metabolic disease, we focused on the simultaneous evaluation of quantitative traits related to liver function and CVD. Using a combination of low-coverage (5×) whole-genome sequencing and targeted genotyping, deep genotype imputation based on the TOPMed reference panel^7^, and genome-wide association study (GWAS) meta-analysis, we analyzed 12 liver-related blood traits (including liver enzymes, blood lipids, and markers of iron metabolism) in up to 203,476 people from three population-based cohorts of different ancestries. We identified 88 likely causal protein-altering variants that were associated with one or more liver-related blood traits. We identified several loss-of-function (LoF) variants reducing low-density lipoprotein cholesterol (LDL-C) or risk of CVD without increased risk of liver disease or diabetes, including variants in known lipid genes (e.g. *APOB*, *LPL*). A novel LoF variant, *ZNF529*:p.K405X, was associated with decreased levels of LDL-C (P=1.3×10^−8^) but demonstrated no association with liver enzymes or non-fasting blood glucose levels. Silencing of *ZNF529* in human hepatocytes resulted in upregulation of LDL receptor (LDLR) and increased LDL-C uptake in the cells, suggesting that inhibition of *ZNF529* or its gene product could be used for treating hypercholesterolemia and hence reduce the risk of CVD. Taken together, we demonstrate that simultaneous consideration of multiple phenotypes and a focus on rare protein-altering variants may identify promising therapeutic targets.

## MAIN TEXT

We combined several approaches for genomic discovery to identify independent variants associated with 12 liver-related phenotypes (see **Supplementary Figure 1** for an overview). The 12 traits we examined were related to: i) blood lipid levels which impact cardiovascular, neurological and eye-related diseases: total cholesterol (TC), low-density lipoprotein cholesterol (LDL-C), high-density lipoprotein cholesterol (HDL-C) and triglyceride (TG) levels; ii) C-reactive protein (CRP; only values <15 mmol/L were included) which is predictive of cardiovascular disease;^8^ iii) enzymes which mainly reflect liver function: alanine aminotransferase (ALT), aspartate aminotransferase (AST), alkaline phosphatase (ALP) and gamma-glutamyltransferase (GGT); and iv) iron-related phenotypes: serum iron, total iron binding capacity (TIBC), and transferrin saturation percentage (TSP).

Using four primary discovery designs, we identified 763 unique variants within 340 genomic regions (i.e. loci) associated with at least one of the 12 quantitative liver-related blood traits. We identified genome-wide significant associations with at least one trait at 88 presumed causal protein-altering variants, of which 9 result in LoF of a specific protein – 3 frameshift indels and 6 premature stop codons.

First, we tested for association with liver-related traits among 69,479 participants (**Supplementary Table 1**) in the population-based Nord-Trøndelag Health Study (HUNT) in Norway.^9^ 26 million polymorphic genetic variants were imputed with sufficient quality and at least 10 alleles using the TOPMed multi-ethnic reference panel consisting of 60,039 deeply sequenced genomes.^7^ We used a linear mixed model^10^ to account for relatedness in the HUNT sample set and identified 246 genome-wide statistically significant (P<5×10^−8^) associations between independent genomic loci and one or more traits (**Supplementary Figure 2** and **Supplementary Table 2**). At 28 of the 246 loci, the most strongly associated variant (i.e. the locus index variant) alters (N=26) or results in LoF (N=2) of the protein. We considered these 28 variants as presumed casually related to the trait of interest.

Second, we conducted step-wise conditional analyses across the 246 primary loci and identified an additional 189 independent associated variants (**Supplementary Table 3**). These include 30 additional protein-altering variants, including 2 LoF variants, that were significantly associated with one or more liver traits but were independent of the original index variant. Notable variants include *TM6SF2*:p.L156P,^11^ associated with TG and ALT (P_TG_=8×10^−9^, P_ALT_=6×10^−10^), and a rare variant in *PCSK9* (*PCSK9*:p.N157K)^12^ associated with LDL-C (beta=-1.1, minor allele frequency [MAF]=0.05%, P=1×10^−14^). We identified significant associations with ALP and four independent coding variants in the alkaline phosphatase gene (*ALPL*:p.R75H, p.M115T, p.E69K, p.E114K), and with the liver trait ALT, we identified association with four independent coding variants in the gene encoding the enzyme alanine aminotransferase 1 (*GPT*: p.A87V, p.G128S, p.P234L, p.V452L). ALP was also associated with five independent coding variants in *GPLD1* (p.N103S, p.P336L, p.R181C, p.S452L, p.V815Sfs*46), encoding the enzyme phosphatidylinositol-glycan-specific phospholipase D,^13^ which degrades a glycosylphosphatidylinositol anchor of proteins to the cell surface and is known to interact with apolipoprotein A-containing complexes.^14^ Interestingly, we also identified two protein-altering variants in *APOE*:p.C130R) demonstrated substantial pleiotropy – showing association with CRP, LDL-C, TC, HDL-C, and ALT;p.R176C was associated with LDL-C, TC and HDL-C. Together, these two variants define the well-known APOE 2, 3, and 4 haplotypes.

Third, we performed association testing for the 12 liver-related blood traits in up to 57,060 HUNT Study participants based on directly genotyped variants that were included as custom content on the array and not part of the primary GWAS (see **Online Methods** for details). This identified 14 additional protein-altering variants, including 4 LoF variants (P<5×10^−8^; **Supplementary Table 4**). Thirteen of these variants were rare (ranging from 1 in 177 to 1 in 6313 individuals). The 14 variants included 8 variants originating from low-coverage (5×) whole-genome sequencing of 2,202 HUNT Study participants. In addition, we identified a significant association with three *LDLR* variants (p.D180N, p.R248W, and p.M484V with 19, 9 and 30 allele copies, respectively). Lastly, we identified three nonsense variants in *APOB* that demonstrated significant association with LDL-C, bringing the total number of independent associated *APOB* variants to seven (p.L12_L14del, p.R1333X, p.K1813X, p.H1923R, p.W3087X, p.R3527Q, p.S4338N). Each of these three HUNT-specific strategies identified novel liver-trait associated variants that would not otherwise be found from imputation-based approaches.

Fourth, we performed a trans-ancestry meta-analysis by combining summary statistics based on the primary discovery effort in HUNT with additional GWAS statistics from Sardinia (SardiNIA cohort)^15^ and Japan (Biobank Japan)^16^, resulting in an analysis of up to 203,476 participants and 31.5 million unique variants (**Supplementary Figure 3**). This analysis identified 388 significant variants, identifying 93 additional loci (**Supplementary Table 5**) and an additional of 16 protein-altering locus index variants, including the previously described Sardinian-specific LoF variant *HBB*:p.Q40X, which has been associated with beta thalassemia^17^ and decreased LDL-C and TC (Table 1).^18^

**Table 1:** Loss-of-function variants associated with liver-related traits

| Gene variant | Chr:pos | Minor/major allele | MAF % (MAC) | Trait | Beta SD | SE | P | Discovery method | Novelty of variant-trait association |
| --- | --- | --- | --- | --- | --- | --- | --- | --- | --- |
| *APOB* p.K1813X | chr2:21011431 | A/T | 0.009 (10) | LDL-C | -2.63 | 0.35 | $1.1 \times 10^{-13}$ | Custom array – predicted nonsense | Novel (in known LDL-C locus; known gene<sup>19,20</sup>) |
| *APOB* p.R1333X | chr2:21013379 | A/G | 0.009 (10) | LDL-C | -2.80 | 0.35 | $2.5 \times 10^{-15}$ | Custom array – predicted nonsense | Novel (in known LDL-C locus; known gene,<sup>19,20</sup> variant previously linked to familial hypobetalipoproteinemia<sup>21</sup>) |
| *APOB* p.W3087X | chr2:21007608 | T/C | 0.011 (12) | LDL-C | -2.79 | 0.32 | $5.1 \times 10^{-18}$ | Custom array – predicted nonsense | Novel (in known LDL-C locus; known gene<sup>19,20</sup>) |
| *GPLD1* p.V815Sfs*46 | chr6:24429112 | C/CT | 0.85 | ALP | -0.87 | 0.039 | $2.2 \times 10^{-107}$ | HUNT locus index variant (imputed from TOPMed) | Novel (in known ALP locus<sup>16</sup>; gene experimentally linked to liver disease<sup>22</sup>) |
| *HBB* p.Q40X | chr11:5226774 | A/G | 4.81 | TC | -0.48 | 0.048 | $5.4 \times 10^{-23}$ | Trans-ancestry meta-analysis index variant | Novel (previously associated with beta thalassemia,<sup>17</sup> which has been related to LDL-C and TC<sup>23</sup>) |
| *LIPC* p.G247Afs*11 | chr15:58545904 | ACG/A | 0.16 (221) | HDL-C | 0.58 | 0.081 | $1.1 \times 10^{-9}$ | HUNT conditional analysis (imputed from TOPMed) | Novel (in known HDL locus;<sup>19,20</sup> variant report in carrier with mixed hyperlipidemia but who also carried APOB variants<sup>24</sup>) |
| *LPL* p.S474X | chr8:19962213 | G/C | 11.9 | TG | -0.17 | 0.0057 | $1.3 \times 10^{-196}$ | Trans-ancestry meta-analysis index variant | Known variant at known locus<sup>19,20</sup> |
| *SLC22A1* p.D426Pfs*28 | chr6:160139865 | C/CTGGTAAGT | 39.3 | LDL-C | -0.05 | 0.0060 | $3.3 \times 10^{-9}$ | HUNT conditional analysis (imputed from TOPMed) | Novel (in known LDL-C locus<sup>19,20</sup>) |
| ZNF529 p.K405X | chr19:36547291 | A/T | 0.099 (110) | LDL-C | -0.60 | 0.11 | 1.3x10<sup>-8</sup> | Custom array –<br>observed in low-pass<br>genomes | Novel |

*Chr: Chromosome, pos: position human genome build hg38, MAF: Minor allele frequency, MAC: minor allele count, SE: standard error of the beta, P: P-value, LDL-C: Low-density lipoprotein cholesterol, ALP: Alkaline phosphatase, TC: Total cholesterol, HDL-C: High-density lipoprotein cholesterol, TG: Triglycerides.*

After combining results across all samples and discovery strategies, we were particularly interested in nine variants which appeared to result in LoF of a gene, including the novel association between *ZNF529*:p.K405X and decreased LDL-C (Table 1). We observed 4 additional LoF variants also resulting in substantially decreased LDL-C (3 nonsense variants in *APOB*, and a common frameshift indel in *SLC22A1*; Table 1). The remaining LoF were associations with other blood lipid traits (*HBB*:p.Q40X with TC in addition to LDL-C, *LPL*:p.S474X with TG, and *LIPC*:p.G247Afs*11 with HDL-C). Of the 9 LoF variants, the five within *APOB, LPL, LIPC*, and *ZNF529* were not even nominally significantly associated (P>0.05) with liver enzymes ALT, AST, ALP or GGT (Figure 2b). This suggests that these genetic variants do not cause liver damage, suggesting these genes may serve as potential drug targets to reduce LDL-C.

We expect that protein-altering variants which are the peaks of association represent functional variants that pinpoint biologically-relevant genes and potential drug targets. We also sought novel genes that decreased cardiovascular risk factors (such as LDL-C), but did not increase risk of liver disease or impact liver enzymes. Thus, we focused on the novel association between *ZNF529*:p.K405X and LDL-C (P=1.3×10 ^−8^) since this variant was not associated with liver enzymes (P=0.4 – 0.9 for all 4 liver enzyme traits, up to N=48,569) or (non-fasting) blood glucose (P=0.93, N=54,093 individuals) in HUNT. To experimentally assess the consequence of LoF of zinc finger 529 (*ZNF529*), we transiently knocked-down expression of *ZNF529* in human hepatoma HepG2 cells using siRNA (90.5% reduction). This resulted in significant upregulation of LDL receptor (LDLR) mRNA (increased 91.5%, P=3×10 ^−6^) and protein (83%, P=0.001) (Figure 1). Using labeled LDL-C, we observed that *ZNF529*-silencing resulted in a marked increase in LDL-C uptake by HepG2 cells and 2.2-fold increase in intracellular cholesterol content (P=0.007). These findings suggest that *ZNF529* is a novel regulator of plasma LDL-C via regulation of hepatic LDLR.

**Figure 1.**
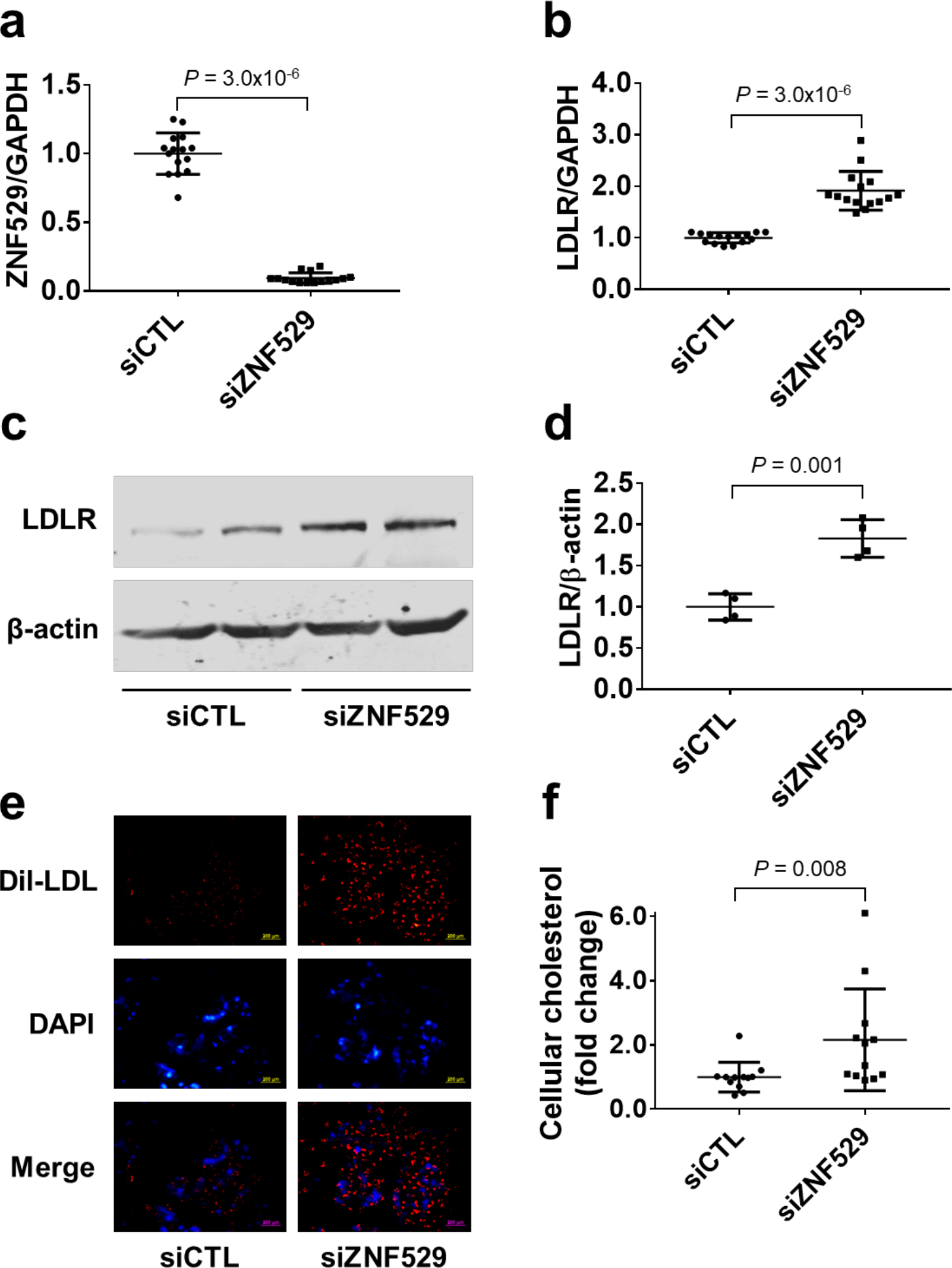
ZNF529 silencing induces LDLR expression and LDL-C uptake. *(a) Efficient silencing of ZNF529 in HepG2 cells via siRNA as shown by qPCR using GAPDH as reference (N=15). **(b)** ZNF529 silencing in HepG2 cells induces LDLR mRNA as shown by qPCR using GAPDH as reference (N=15), **(c, d)** and LDLR protein as shown by Western blot using β-actin as loading control (N=4). **(e)** ZNF529 silencing in HepG2 cells increases LDL-C uptake as evidenced by enhanced fluorescence of Dil-LDL. scale bars=200 µm, (N=6), **(f)** and leads to increased intracellular cholesterol (N=12). Values are presented as mean ± SD showing all points*.

Individuals heterozygous for *ZNF529*:p.K405X (N=109) had a mean LDL-C level of 2.58 mmol/L vs. 3.44 mmol/L in non-carriers. This reduction in LDL-C of 25% in heterozygous carriers is in the range of what is seen for 5 years of treatment with 40mg of statin (−35% change).^25^ We only observed one homozygous female carrier. Despite being obese and server hypertensive, she was alive at age >90 years and had no diagnosis of cardiovascular disease, liver disease, or diabetes, and had an LDL-C level slightly below average for her age group (3.45 mmol/L vs. mean 3.8 mmol/L for women >90 years old). This one individual with a natural absence of both copies of *ZNF529* suggests that homozygous knockout of this gene is compatible with survival.

We also highlight 17 protein-altering variants with an impact >1 standard deviation on the trait (Figure 2a, **Supplementary Figure 4**, and **Supplementary Table 6**). For lipids, protein-altering variants in *APOB, LDLR*, and *PCSK9* that impact LDL-C, and in *CETP* that impact HDL-C, are well known.^19^ However, for liver enzyme traits, *TNK1*:p.G574V is a new finding to complement genes previously known to impact liver enzymes including *ALPL* (with ALP),^26^ *GPLD1* (with ALP), and *GPT* (with ALT).^27^ This rare *TNK1* variant, present in 46 individuals (1 in 814 individuals), was identified in Norwegian sequenced samples, genotyped using the custom array, and observed to have a large impact on ALP (beta=1.2, P=1.7×10^−14^).

To further characterize genes that may be involved in these liver-related phenotypes, we performed gene-based burden tests, using SKAT-O as implemented in SAIGE-GENE,^28^ for all protein-altering variants with frequency below 0.5% in the HUNT dataset. Although twenty-eight unique genes were significantly associated (P<2×10^−7^) with at least one of 12 liver traits (**Supplementary Table 7**), in only two cases was the gene-based evidence for association substantially stronger than the gene-based test relative to the strongest single variant: rs147998249 with the gene *GPT* associated with ALT (P=2.35×10^−60^); rs138587317 with the gene *ALPL* associated with ALP (P=2.9×10^−239^). These data suggest there are multiple, functional coding rare variants in each of these two genes as noted above. Gene-based burden results, which are independent of nearby signals, may point to the functional gene. We see this at well-known genes *CETP* and *ABCA1* for HDL-C; *PCSK9, LDLR* and *APOB* for LDL-C; the *CRP* gene for CRP, *TF* for TIBC; and *GPLD1* for ALP (**Supplementary Table 7**). Additionally, burden tests for LoF of *APOB* variants indicated no association with liver enzymes for heterozygous LoF carriers (**Supplementary Table 8**).

**Figure 2.**
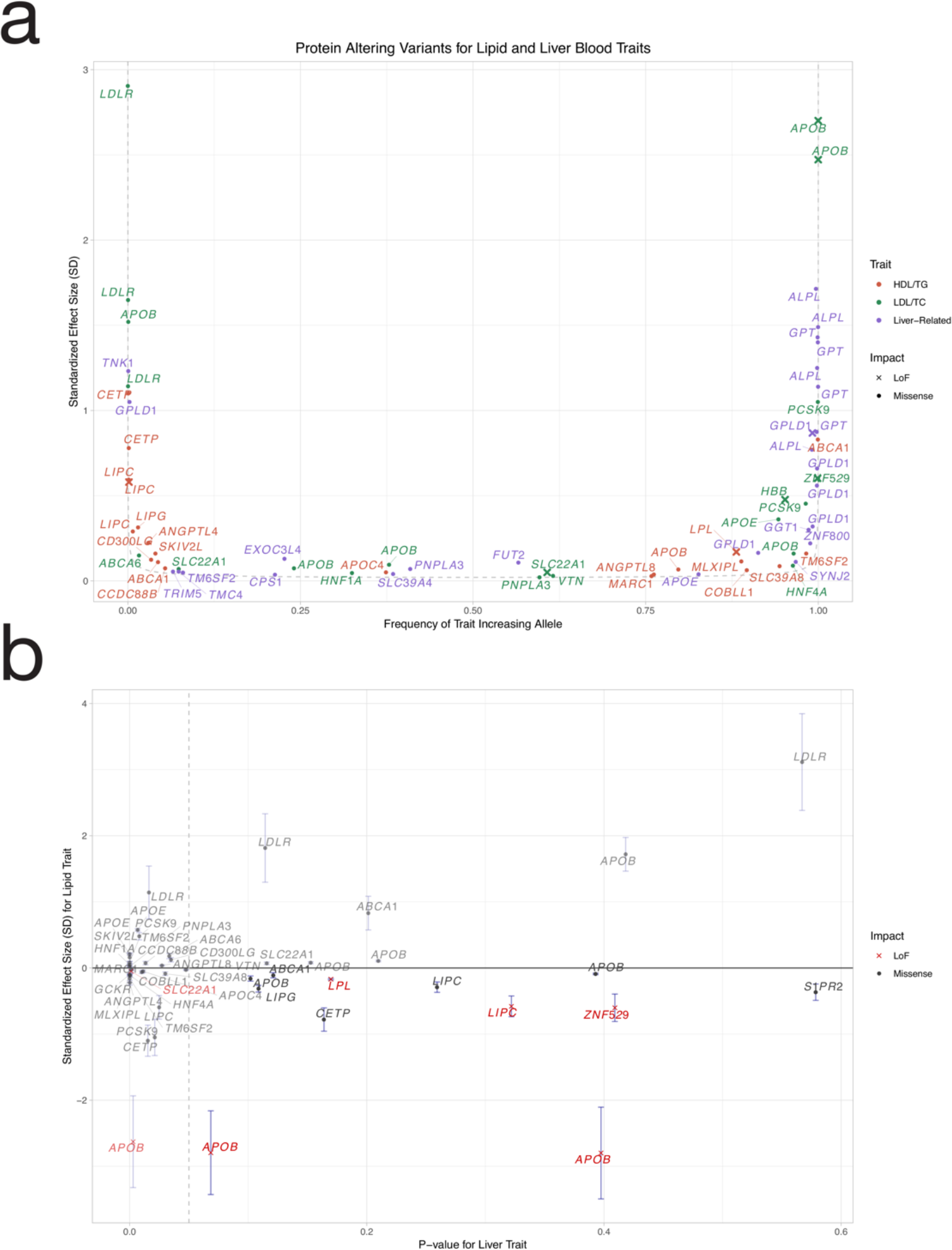
Protein Altering Variants for Lipid and Liver Blood Traits. **(a)** Smile plot comparing the frequency of the blood-trait increasing allele with the allele’s effect size for protein altering variants significantly (P<5×10^−8^) associated with a lipid (HDL-C, LDL-C, TG, TC) or liver (ALT, ALP, AST, GGT) trait. The most significant variant is shown for variants with significant association for multiple traits. Color indicates the trait category for which the variant is significant, with loss-of-function variants shown as x. Power curve (dashed line) denotes estimated 90% power in the meta-analysis with a sample size of N=210,000 at alpha=5×10^−8^. **(b)** For any variant significantly (P<5×10^−8^) associated with a lipid trait (HDL-C, LDL-C, TG, TC), the maximum effect size in terms of the allele associated with good lipid health (e.g. lowered LDL-C, increased HDL-C, lowered TG, and lowered TC) is compared to the minimum p-value for association with liver trait (ALT, ALP, AST, GGT). Nominal P-value of 0.05 (vertical dashed line) is indicated to highlight variants in the bottom right quadrant which lack significance for association with liver traits. These variants are better drug target candidates given estimated favorable lipid-effects on health and absence of association with potentially unfavorable liver traits.

To expand our understanding of the 88 protein-altering variants, to investigate their impact on disease, and to evaluate potential consequences of targeting the implicated gene or its gene product, we imputed the TOPMed reference panel^7^ into the UK Biobank and performed a phenome-wide association studies (PheWAS) across 1,342 ICD code-defined disease groups,^29,30^ in 408,961 people of white British ancestry. 77 of the 88 protein-altering variants could be imputed sufficiently well (R^2^>0.3). For 29 variants, we found a phenome-wide significant (P<3.5×10^−5^) association with one or more diseases (Figure 3, **Supplementary Figure 5**, and **Supplementary Table 9**).

**Figure 3:**
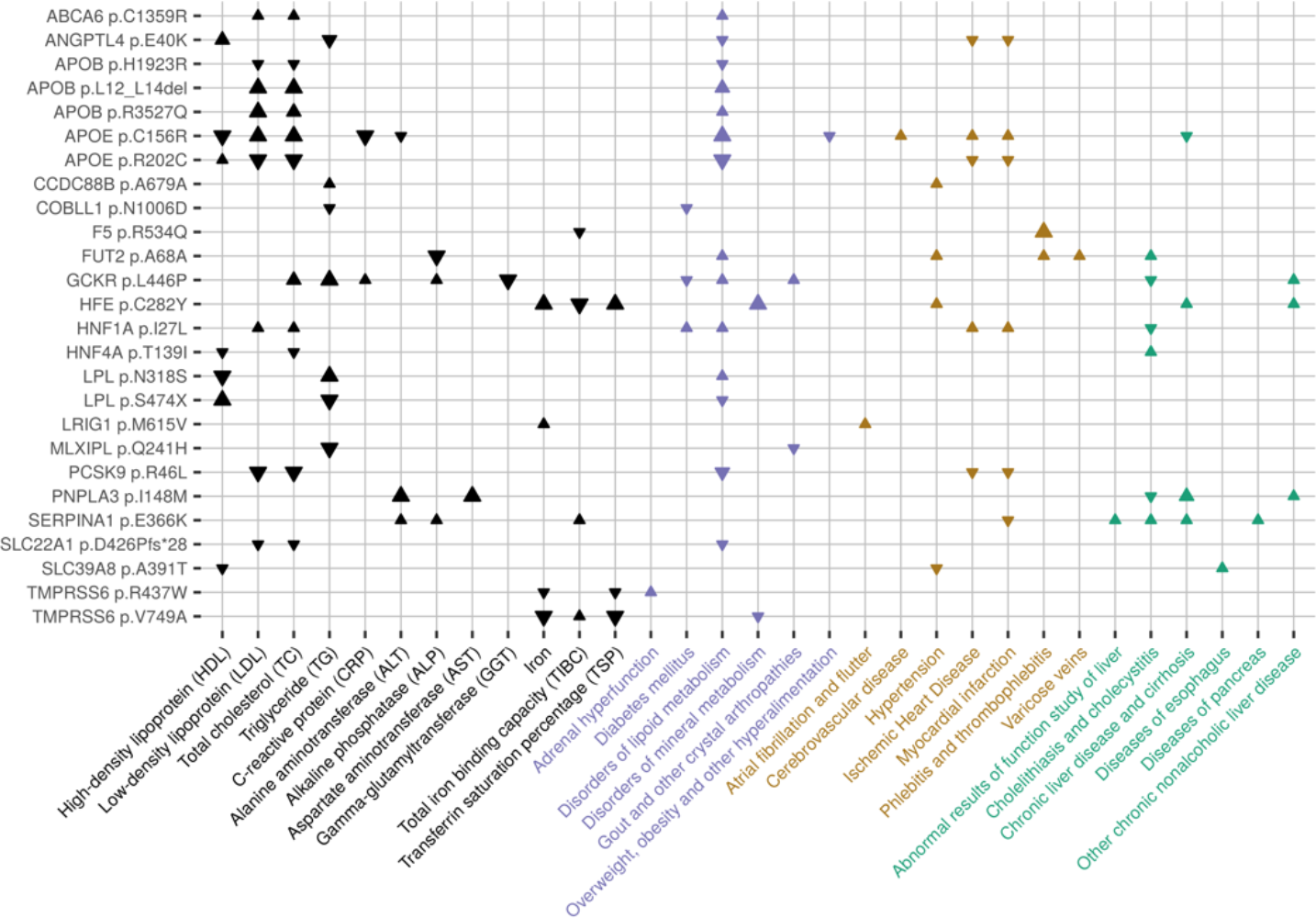
Phenome-wide associations in UK Biobank (N=408,961 participants) based on protein-altering variants with impact on liver-related blood traits in The HUNT Study (N=69,479). 29 protein-altering variants associated with liver-related traits had additional associations with selected cardiovascular, liver, and metabolic phenotypes. Arrows denote the direction of effect for the minor allele. Larger arrows signify more significant associations. Please see Supplementary Figure 5 for the full phenome-scan across all traits and variants. The ZNF529 LoF variant could not be evaluated in the UK Biobank.

To identify potentially useful drug targets that may reduce blood lipid levels and risk of coronary artery disease (CAD) and type 2 diabetes (T2D) without an increased risk of fatty liver disease, we attempted to identify variants that decreased LDL-C and/or TG, but were not associated with changes in liver enzyme levels (P>0.05, Figure 3, **Supplementary Figure 5** and **Supplementary Tables 6-8**), suggesting that liver function was not altered. The four variants with this pattern of association include *COBLL1* p.N497D which is associated with decreased TG levels and decreases risk of T2D in both HUNT and UK Biobank and of liver disease in HUNT. HDL-C-associated *ANGPTL4* p.E40K appears to decrease risk of T2D, CAD and hypertension whereas *LPL* p.N318S appears to decrease risk of CAD, liver disease and hypertension. *APOB* LoF variants decrease LDL-C and risk of CAD, but may also have a beneficial impact on CAD, T2D and hypertension (Figure 3, **Supplementary Figure 5** and **Supplementary Tables 6-8**).

From the PheWAS, we observed other interesting associations. For example, we identify alleles that reduce risk of myocardial infarction (MI) but increase risk liver and pulmonary disease for *SERPINA1*:p.E366K, which is known for causing alpha-1-antitrypsin deficiency^31^. This pattern suggests caution in ongoing efforts to treat acute MI with exogenous administration of alpha-1-antitrypsin.^32,33^ Another interesting and novel finding from UK Biobank PheWAS is the association between *TMPRSS6*:p.V727A and an increased risk of ‘nonspecific chest pain’. We initially found this allele to be associated with decreased iron and TSP in HUNT, – perhaps suggesting that the association with chest pain might be explained by anemia-induced cardiac ischemia (Figure 3, **Supplementary Figure 5** and **Supplementary Tables 7, 9**). Further studies are obviously warranted to uncover the biological mechanisms underlying these associations, however, each of them could help inform clinical implications of targeting the underlying gene or gene product.

In summary, by using four complementary approaches for genomic discovery: sequencing, imputation, array-based genotyping and trans-ancestry meta-analysis, we identified >300 loci associated with liver-related quantitative traits, including 88 presumed causal protein-altering variants. By considering disease end-point associations with disease phenotypes for protein-altering variants, we prioritize several novel genes as potential drug targets. The newly uncovered association and *in vitro* studies indicate that *ZNF529* LoF is associated with lower plasma LDL-C which could be explained by induction of LDLR in hepatic cells and increased LDL-C uptake. While these findings indicate a therapeutic potential for lowering plasma LDL-C by ZNF529 inhibition, further studies are warranted to elucidate the mechanisms by which ZNF529 regulates LDLR and LDL-C uptake in the liver. We further confirmed several genes as promising drug targets for cardiometabolic disease with no expected impact on liver disease by testing association with impactful protein-altering variants in *APOB* and *LPL*, to reduce LDL-C and risk of coronary artery disease, and *COBLL1*, to reduce TG and risk of type 2 diabetes. Future functional studies to rule out their role in liver and other cardiometabolic disease will likely contribute to better prediction of outcomes and disease progression and facilitate development of personalized treatments.

All together, we demonstrate that identifying rare protein-altering variants and careful consideration of multiple phenotypes in well-powered studies may point to promising drug targets. We used a variety of approaches to identify rare protein-altering variants, and we found that if exome sequencing is prohibitively expensive, sequencing a subset of samples followed up with a custom genotyping array can be a viable strategy to identify impactful rare variants.

## Supporting information

Supplementary Figurs

## URLs

SAIGE/SAIGE-GENE [https://github.com/weizhouUMICH/SAIGE] EPACTS [https://github.com/statgen/EPACTS] liftOver: [https://genome.ucsc.edu/cgi-bin/hgLiftOver]

## ACKNOWLEDGEMENTS

The Nord-Trøndelag Health Study (The HUNT Study) is a collaboration between HUNT Research Centre (Faculty of Medicine and Health Sciences, NTNU, Norwegian University of Science and Technology), Nord-Trøndelag County Council, Central Norway Regional Health Authority, and the Norwegian Institute of Public Health. The K.G. Jebesen Center for Genetic Epidemiology is financed by Stiftelsen Kristian Gerhard Jebsen; Faculty of Medicine and Health Sciences, NTNU, Norwegian University of Science and Technology (NTNU) and Central Norway Regional Health Authority. Whole genome sequencing for the HUNT study was funded by HL109946. Whole genome sequencing (WGS) for the Trans-Omics in Precision Medicine (TOPMed) program was supported by the National Heart, Lung and Blood Institute (NHLBI). WGS for NHLBI TOPMed studies (Freeze 5: AACAC, AFGen, Amish, ARIC+VTE, Asthma_Afr, Asthma_CR, CHS, COPDGene, Framingham, GeneStar, GENOA, GenSalt, GOLDN, HyperGen, Jackson, MESA, MLOF/Hemophilia, OMG-SCD, PharmHU, REDS-III-SCD, SAFHS, Sarcoidosis, SARP, THRV, walk-PhaSST, WHI) was performed at Baylor Human Genome Sequencing Center, Broad Institute of MIT and Harvard, Illumina Genomic Services, Macrogen Corp, New York Genome Center, University of Washington Northwest Genomics Center). Centralized read mapping and genotype calling, along with variant quality metrics and filtering were provided by the TOPMed Informatics Research Center (3R01HL-117626-02S1). Phenotype harmonization, data management, sample-identity QC, and general study coordination, were provided by the TOPMed Data Coordinating Center (3R01HL-120393-02S1). We gratefully acknowledge the studies and participants who provided biological samples and data for TOPMed. This research has been conducted using the UK Biobank Resource under application number 24460. J.B.N. is supported by grants from the Danish Heart Foundation (16-R107-A6779) and the Lundbeck Foundation (R220-2016-1434). O.R. was supported by the American Heart Association Postdoctoral Fellowship 19POST34380224 and the Michigan-Israel Partnership Research Grant. S.A.L. is supported by NIH grant 1R01HL139731 and American Heart Association 18SFRN34250007. I.S. is supported by a Precision Health Scholars Award from the University of Michigan Medical School. L.V. was supported by NIH grant R01-HL123333. P.N. was supported by NIH grant K08-HL140203. J.Z. was supported by NIH grant R01-HL138139. Y.E.C was supported by NIH grants R01-HL068878 and R01-HL137214. C.J.W. was supported by NIH grants R35-HL135824, R01-HL127564, R01-HL117626-02-S1, and R01-HL130705. The SardiNIA research was supported in part by the Intramural Research Program of the National Institute on Aging, National Institutes of Health (NIH) (contracts N01-AG-1-2109 and HHSN271201100005C). This work was also supported by the National Institutes of Health (NHLBI grant HL117626).

## AUTHOR CONTRIBUTIONS

J.B.N, S.E.G, I.S, W.Z., L.F., and S.A.G.T. performed the computational analyses. O.R., Y.L., L.V. and J.Z. performed wet lab experiments. J.B.N., K.H., Y.E.C., and C.J.W. designed and supervised the study. All authors contributed to manuscript preparation and read, commented on and approved the manuscript.

## DISCLOSURES

GRA works for Regeneron Pharmaceuticals. PN reports grant support from Amgen, Apple, and Boston Scientific, and is a scientific advisor to Apple. SAL receives sponsored research support from Bristol Myers Squibb / Pfizer, Bayer HealthCare, and Boehringer Ingelheim, and has consulted for Bristol Myers Squibb / Pfizer. PTE has consulted for Novartis, Quest Diagnostics and Bayer AG. STW has received royalties from UpToDate. S.A. holds equity in 23andMe, Inc.

## ONLINE METHODS

### Discovery cohorts

#### HUNT

The Nord-Trøndelag Health Study (HUNT) is a population-based health survey conducted in the county of Nord-Trøndelag, Norway, since 1984.^9^ Participation in the HUNT Study is based on informed consent and the study has been approved by the Data Inspectorate and the Regional Ethics Committee for Medical Research in Norway (REK: 2014/144). We included a total of 69,479 individuals with values for at least one of the traits examined (ALT, ALP, AST, CRP, GGT, HDL-C, LDL-C, TC, TG, Iron, TIBC, TSP). Genotyping was performed using the Illumina Human CoreExome v1.1 array with 70,000 additional custom content beads.^34,35^ Variants were selected for genotyping if they were: protein-altering (N=13,618); modestly associated with lipids in HUNT but not tested in large consortia (N=960); identified as causing familial hypercholesterolemia in Norwegian patients (N=110); or predicted to result in a loss-of-function of one of the 56 ACMG genes (N=27,144, **Supplementary Table 10**). Additionally, we selected missense variants with 2 or more copies (N=8,720) and nonsense variants with 1 or more copy (N=756) identified from low-pass sequencing of 2,202 HUNT samples.

Imputation was performed from 60,039 TOPMed reference genomes using Minimac3 and variants with imputation quality >0.3 were retained. To account for relatedness within the sample, we performed association testing using the linear mixed model with genetic relationship matrix as implemented in SAIGE.^10^ Conditional analysis was performed with the same analysis tools and command line options as the association analysis. Conditional analysis was performed by adding the lead-SNP(s) in a step-wise manner as covariate(s) into the SAIGE step1 parameter estimation until the variant with smallest P-value in the locus was >5×10^−8^.

#### Biobank Japan

Biobank Japan (BBJ) is a multi-institutional hospital-based registry of ~200,000 individuals from 66 Japanese hospitals collected from 2003-2007. Genotype, imputation, and QC were performed as described previously.^16^ Briefly, samples were genotyped with Illumina HumanOmniExpressExome or a combination of the Illumina HumnOmniExpress and HumanExome BeadChips and imputed using 1000 Genomes Project Phase 1 version 3 East Asian reference haplotypes. Publicly available summary statistics from linear regression assuming an additive model for quantitative measures of ALP, ALT, AST, CRP, GGT, HDL-C, LDL-C, TC, and TG were used. Quantitative traits were adjusted for age, sex, top 10 PCs of genetic ancestry, and disease status for 47 target diseases. Sample sizes for traits ranged from 70,567 to 134,182.^16^

#### SardiNIA

6,602 individuals from four villages in the Lanusei valley on Sardinia (>60% of the adult population) were genotyped on four different Illumina Infinium arrays: OmniExpress, Cardio-Metabochip,^36^ Immunochip,^37^ and Exome Chip. Low-depth (~4× coverage) whole-genome sequencing on 3,839 individuals was performed, of which 2,340 were also genotyped. Imputation of 1.1m indels and 24.1m biallelic single nucleotide variants was performed using Minimac3^38^ and markers with imputation quality >0.3 (or >0.6 if MAF<1%) were retained. Samples, genotyping, sequencing and variant calling have been previously described.^18^

Liver traits (ALT, AST, CRP, GGT, HDL-C, Iron, LDL-C, TC, TG and transferrin) from the first visit were measured except LDL-C, which was computed using the Friedewald Equation.^15^ Association analyses were performed for liver traits assessed in 5,570 – 5,942 individuals (median=5,917) using the age, age^2^ and sex-adjusted inverse-normalized residuals of the outcomes as input to the Efficient Mixed Model Association eXpedited (EMMAX)^39^ single variant test (i.e. a linear model with a kinship matrix) as implemented in EPACTS. Genomic control correction was not applied as the lambda values were not inflated (range 0.97 to 1.02, **Supplementary Table 11**).

### Meta-analyses

As the summary statistics from SardiNIA and Biobank Japan were in Human Genome Build hg19, the positions were mapped to Human Genome Build hg38 using liftOver. The genomic control corrected summary statistics from the contributing cohorts were combined with METAL^40^ using inverse variance weighted meta-analysis. Meta-analysis included SardiNIA, Biobank Japan, and HUNT for all traits, with the exception of ALP, Iron, and TIBC which were only available from HUNT and Biobank Japan. TSP was available only in HUNT and not meta-analyzed.

### PheWAS in UK Biobank

Association results for 1,342 trait groups (PheCodes)^29^ in UK Biobank were generated using SAIGE.^10^ Phenotypes were grouped by combining ICD-9 and ICD-10 codes of closely related traits following previously published methods.^30^ Analysis was performed on the white British subset of UK Biobank after imputation with the TOPMed reference panel. Sex, birth year, and 4 principle components were included as covariates. Significance was determined based on Bonferroni correction for the number of traits tested (P<3.5×10^−5^).

### Gene-based SKAT-O tests

The exome-wide gene-based SKAT-O tests were performed using SAIGE-GENE v36^28^ for all 12 liver traits based on the TOPMed-imputed HUNT data. Missense and stop-gain variants annotated by ANNOVAR^41^ with MAF≤0.005 are included. Conditional analysis was performed to condition on the most significant single variant association signal within 500 kB of the gene. We selected a significance threshold of 2.5×10^−6^ accounting for 20,000 genes and 12 traits.

### Replication attempt of ZNF529:p.K405X in MGI

The Michigan Genomics Initiative (MGI) is a repository of electronic medical record and genetic data at Michigan Medicine (N~58,000 participants). MGI participants were enrolled during pre-surgical encounters at Michigan Medicine and provided consent to study genetic and electronic health record data for research. The MGI study was approved by the Institutional Review Board of the University of Michigan Medical School. DNA was extracted from blood samples and participants were genotyped using Illumina Infinium CoreExome-24 bead arrays, which includes the same custom content as the HUNT study. Genotype data was imputed to the Haplotype Reference Consortium using the Michigan Imputation Server, providing 17 million imputed variants after standard quality control and filtering. Only European individuals were used for analysis. We attempted to replicate the association with *ZNF529:p.K405X* in 13,319 MGI participants with LDL-C measurements, however, only 1 participant was heterozygous for ZNF529 so the power to detect association was near zero. In contrast, we identified 110 heterozygous individuals in the HUNT discovery study.

### In vitro studies

#### Cells

The HepG2 human hepatoma cell line was obtained from the American Type Culture Collection (ATCC) and cultured at 37°C and 5% CO_2_ in Dulbecco’s Modified Eagle Medium (DMEM, Gibco) supplemented with 10% fetal bovine serum (FBS, Sigma-Aldrich) and 1% Penicillin-Streptomycin (Pen-Strep, Gibco).

#### ZNF529 gene silencing using small interfering RNA (siRNA)

siRNA targeting zinc finger protein 529 (siZNF529: GGCUUUUGGAGUAUGUAGAtt) and non-targeting siRNA control (siCTL) were obtained from Ambion (siRNA IDs s33654 and AM4611, respectively). HepG2 cells were transfected with 20 nM of siZNF529 or siCTL using Lipofectamine RNAiMAX (Invitrogen) in Opti-MEM reduced-serum medium (Gibco) in accordance with the manufacturer’s protocol.^42^ Cellular lipid or protein extraction, RNA isolation or LDL-C uptake assays were conducted 48 h post transfection.

#### Lipid extraction and cholesterol quantification

The lipids of HepG2 cells were extracted using hexane (≥99%, 32293, Sigma-Aldrich) and isopropanol (≥99.5%, A426-4, Fisher Chemicals) at a 3:2 ratio (*v:v*), and the hexane phase was left to evaporate for 48 h. The remaining cells in the plates were disrupted in 0.1 M NaOH for 24 h and an aliquot was taken for measurement of cellular protein using the Bradford protein assay (Bio-Rad). The content of cellular cholesterol was determined spectrophotometrically using a commercially available kit (Wako Chemicals, 999-02601). Cholesterol data was normalized to cellular protein levels.^43,44^

#### RNA isolation, RT-PCR and qPCR

Total RNA was purified from HepG2 cells using the QIAGEN’s RNeasy kit (QIAGEN). cDNA was synthesized using SuperScript III (Invitrogen), and qPCR was performed using SYBR green reagents (Bio-Rad). Gene expression is presented as fold increase compared with RNA isolated from control cells by the comparative CT (2^−ΔΔCT^) method using GAPDH as the reference gene.^43,45^ Primer pairs used for qPCR were obtained from Integrated DNA Technologies and are available in **Supplementary Table 12**.

#### Protein extraction and Western blot

Cells were lysed in radioimmunoprecipitation assay lysis buffer (RIPA buffer, Thermo Scientific) supplemented with a protease inhibitor cocktail (Roche Applied Science). Proteins were resolved in 8% sodium dodecyl sulfate polyacrylamide gel electrophoresis (SDS-PAGE) and transferred to nitrocellulose membranes (Bio-Rad). The membranes were blocked for 1 h at room temperature in tris-buffered saline-Tween 20 (TBST) containing 5% fat-free milk and incubated with primary antibody at 4°C overnight. The following primary antibodies were used: rabbit monoclonal anti-LDLR antibody (Abcam, ab52818, working dilution 1:1000) and mouse monoclonal anti-β-actin antibody (Cell signaling, 8H10D10, working dilution 1:2000). After TBST washing, membranes were incubated with secondary antibodies (LI-COR Biotechnology, donkey anti-rabbit IRDye 926-32212 and donkey anti-mouse IRDye 926-68072, working dilution 1:10000) for 1 hour at room temperature. After TBST washing, bands were visualized and quantified using an Odyssey Infrared Imaging System (LI-COR Biosciences, version 2.1).^43^

#### DiI-LDL uptake assay

3,3’-dioctadecylindocarbocyanine-low density lipoprotein (DiI-LDL, Alfa Aesar) was used to evaluate the cellular uptake of LDL-C^46^ in accordance with the manufacturer’s instructions. Briefly, 48 h following siRNA transfection, HepG2 cells were washed with PBS (x2) and changed to serum-free media supplemented with 0.1% bovine serum albumin (BSA, Sigma-Aldrich). Then, cells were incubated with 10 μg/ml of DiI-LDL for 5 h at 37°C in the dark. Nuclei were stained with 4ʹ,6-diamidino-2-phenylindole (DAPI). After incubation, the cells were washed twice with serum- and probe-free medium. Finally, the cells were visualized using a fluorescent microscope (Olympus, IX71). For each experiment two random fields were chosen and photographed in a blinded fashion. Dil-LDL and DAPI images were merged using ImageJ software (NIH).

#### Statistical analyses for in vitro studies

Statistical analyses were performed using SPSS 24.0 software (SPSS Inc. IBM). Unless indicated otherwise, values are presented as mean ± SD showing all points. All data were tested for normality and equal variance. If the data passed those tests, Student *t* test was used for comparisons between the two groups. If the data did not pass those tests, nonparametric Mann-Whitney *U* test was used. P<0.05 was considered statistically significant.

## DATA AVAILABILITY

Summary statistics will be deposited upon manuscript acceptance.

