## Supplementary Figurs for "Loss-of-function genomic variants with impact on liver-related blood traits highlight potential therapeutic targets for cardiovascular disease"

Table of Contents

Supplementary Figure 1 – Study Design

Supplementary Figure 2 – QQ plots from HUNT analysis

Supplementary Figure 3 – QQ plots from trans-ancestry meta-analysis

Supplementary Figure 4 – Effect size vs. allele frequency for CRP and Iron-related traits

Supplementary Figure 5 – PheWAS plot (full)

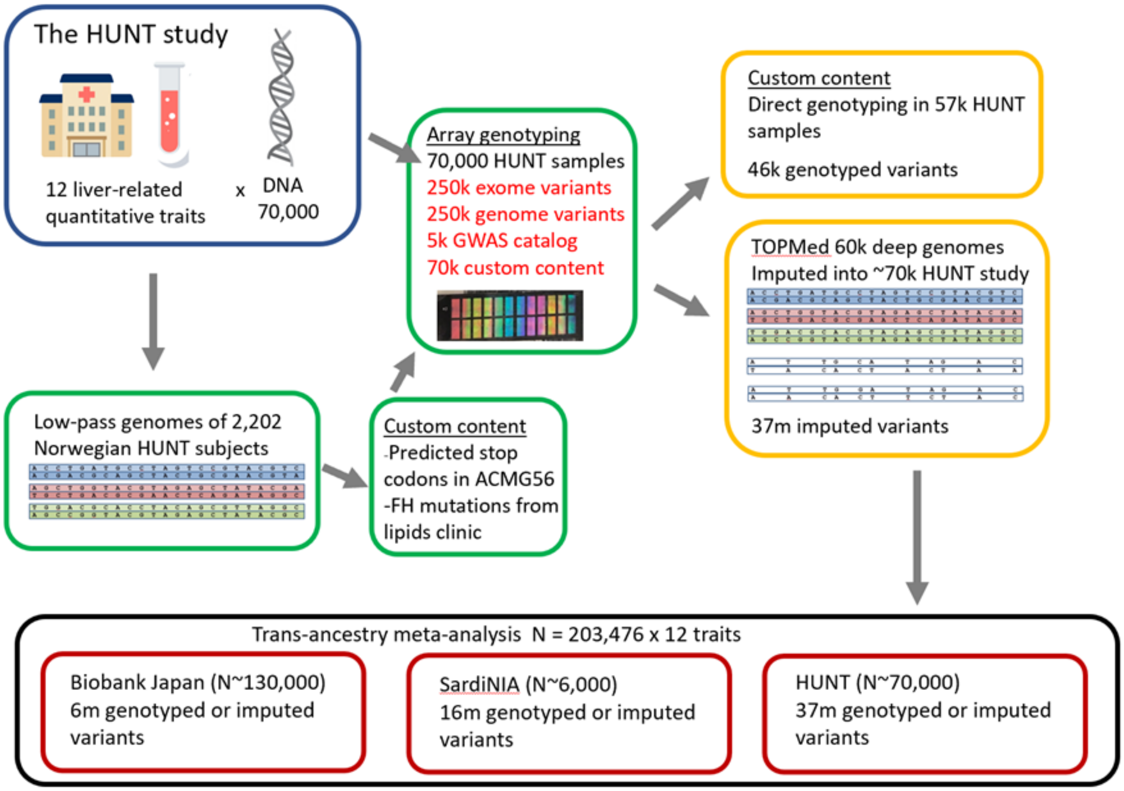


Supplementary Figure 1 – Study Design

Approximately 70,000 individuals from the HUNT study have been genotyped and have available biomarker data. Included in the genotyping array for these individuals were custom content variants selected based on variants identified from whole-genome sequencing of Norwegians, known variants associated with lipid phenotypes, and predicted stop codons in ACMG genes. We analyzed the genotyped variants directly for association with 12 liver-related phenotypes, analyzed the TOPMed imputed results, and performed trans-ethnic meta-analysis with two additional cohorts.

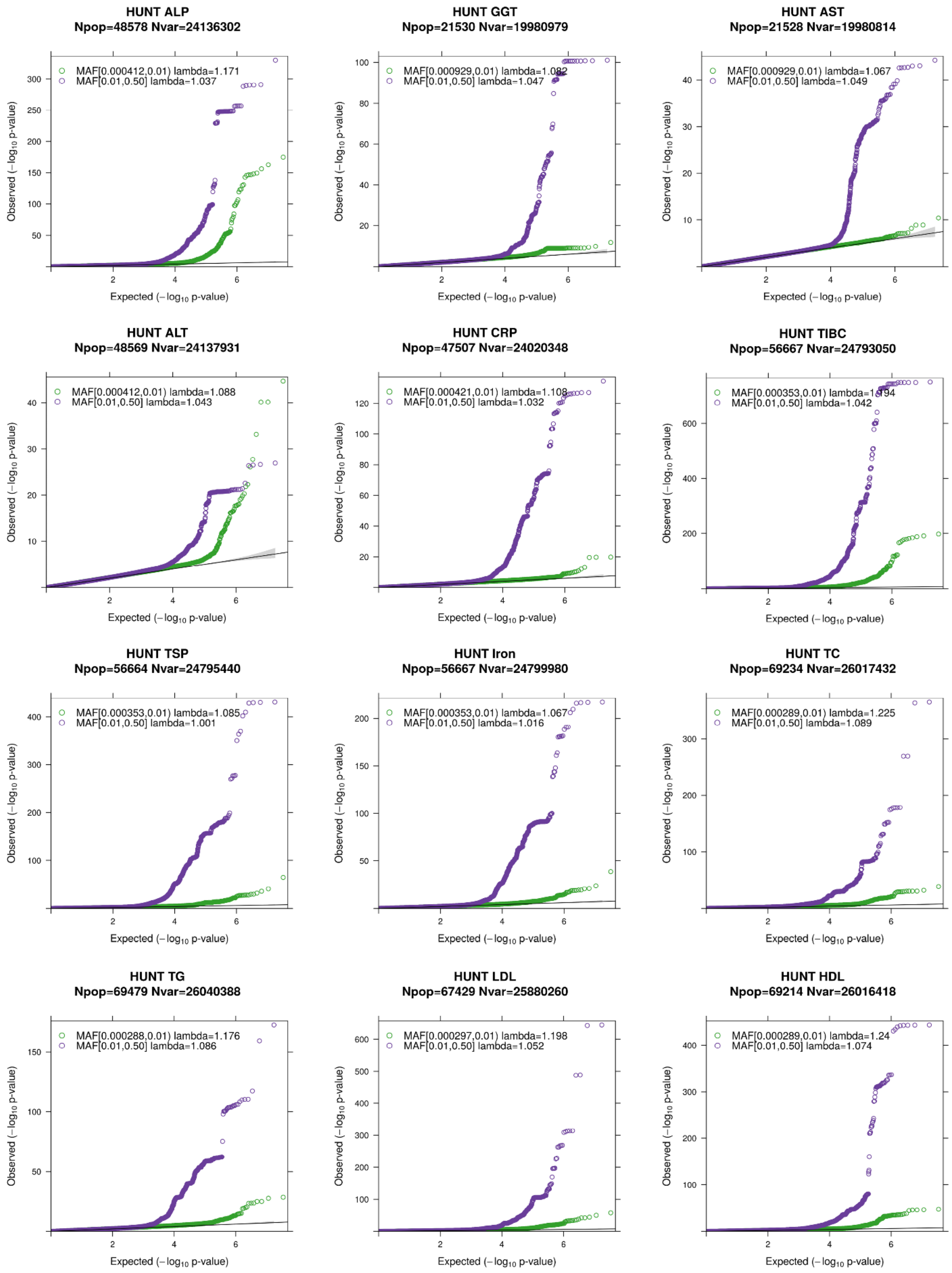


Supplementary Figure 2 – QQ plots from HUNT analysis

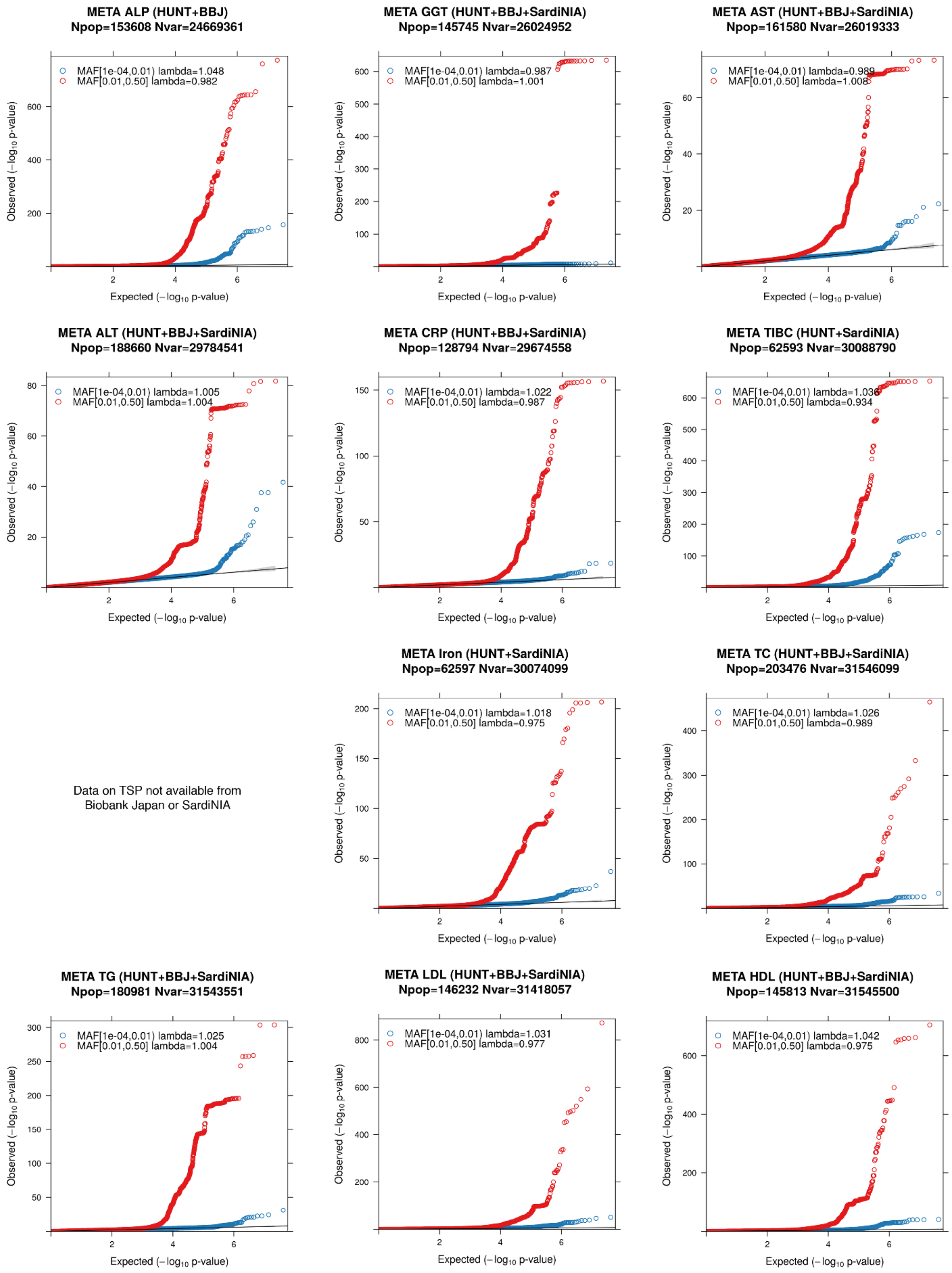


Supplementary Figure 3 – QQ plots from meta-analysis

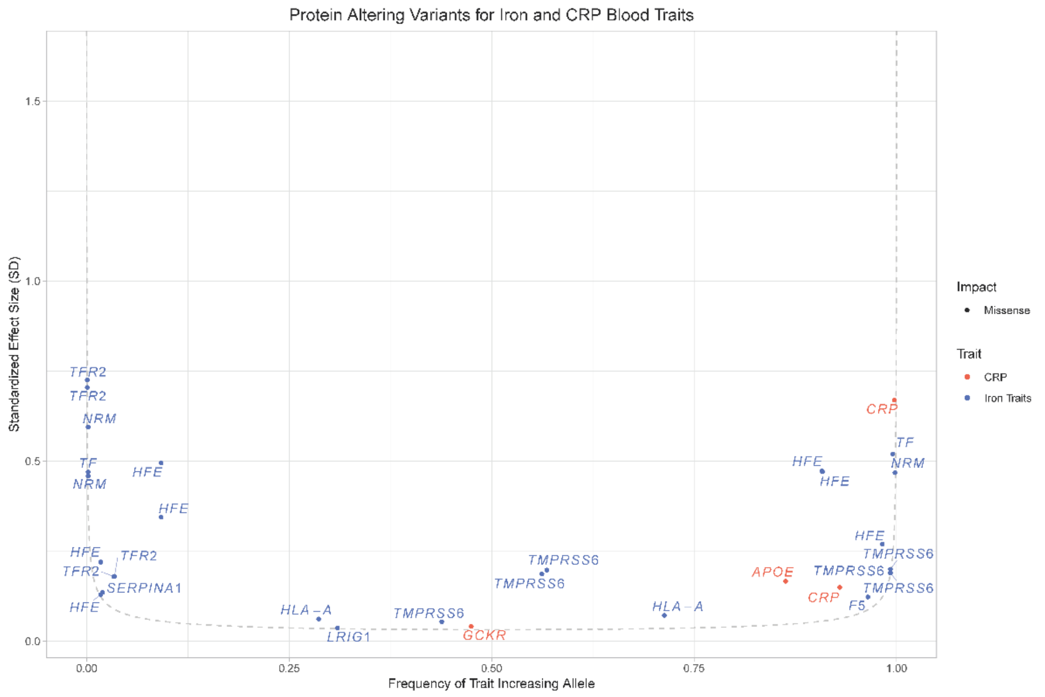


Supplementary Figure 4 - Effect size vs. allele frequency for CRP and Iron-related traits

Smile plot showing protein-altering variants associated with Iron related traits (TIBC, TSP, Iron) and CRP

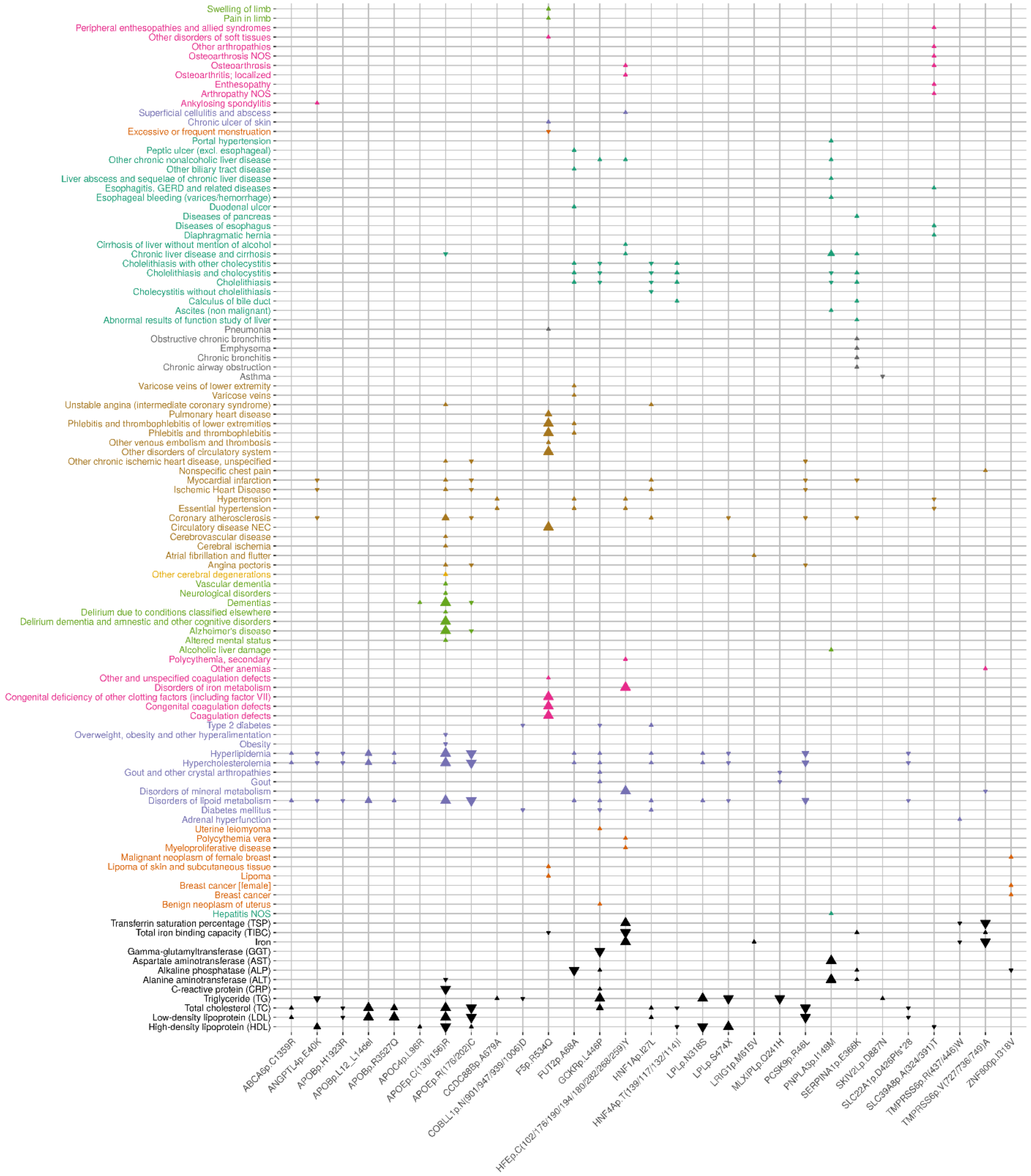


Supplementary Figure 5 – UK Biobank PheWAS results for all protein-altering variants

29 protein altering variants were significantly (P-value < $3.5x10^{-5}$) associated with additional phenotypes within UK Biobank. Arrows denote the direction of effect for the minor allele. Larger arrows signify more significant associations.
